# Silk-based antimicrobial peptide mixed with recombinant spidroin creates functionalized spider silk

**DOI:** 10.1101/2021.03.26.437269

**Authors:** Frank Y.C. Liu

## Abstract

Surgical site infection (SSI) from sutures is a global health emergency because of the antibiotic crisis. Methicillin-resistant S. aureus and other emerging strains are difficult to treat with antibiotics, so drug-free sutures with antimicrobial properties are a solution. Functionalized spider silk protein (spidroin) is a candidate for its extraordinary strength because it has a large repetitive region (150Rep) that forms crosslinked beta-sheets. The antimicrobial peptide HNP-1 can be connected to recombinant spidroin to create antimicrobial silk. Ni-NTA purified 2Rep-HNP1 fusion protein was mixed with recombinant NT2RepCT spidroin at 1:25, 1:20, 1:10 ratios, and spun into silk fibers by syringe-pumping protein into a 100% isopropanol bath. Beta-sheet crosslinking of the identical 2Rep regions tagged the 2Rep-HNP1 permanently onto the resultant silk. Silk showed no sign of degradation in an autoclave, PBS, or EtOH. The tagged 2Rep-HNP1 retained broad-spectrum antimicrobial activity >90% against S. aureus and E. coli as measured by log reduction and radial diffusion assay. Furthermore, a modified expression protocol increased protein yield of NT2RepCT 2.8-fold, and variable testing of the spinning process demonstrated the industrial viability of silk production. We present a promising suture alternative in antimicrobial recombinant spider silk.

## 2. Introduction

Surgical site infections (SSI) are a serious health problem worldwide, representing more than 20% of the 4.1 million cases of healthcare-associated infections in the EU each year.[1] Staphylococcus aureus, specifically the methicillin-resistant strain (MRSA), is now the leading cause of SSI in the US.[2] This poses a significant economic burden to patient hospital care, averaging $25k to patients in the US.[3, 4] SSI often come through contamination of sutures and other equipment, because they can act as a reservoir for infection.[5] The traditional strategy to prevent SSI is to inject antibiotics to the surgery site, and newer methods improve on this by incorporating coatings on sutures, but all still rely on antibiotics.[5–7] The excessive use of antibiotics to treat microbial infections, however, has led to resistant strains that reduce the effectiveness of treatments, leading to the antibiotic crisis.[8] As such, there is a dire need to research alternatives to antibiotic-based sutures that are less likely to develop bacteria resistance in order to decrease SSI.[5, 9–11]

Antimicrobial peptides, polycationic polymers, silver ions, and have been shown to be effective coatings.[12–14] Antimicrobial peptides (AMP) display broad-spectrum antimicrobial activity regardless of antibiotic resistance, and have been shown to display antibiofilm properties as well, making them a good candidate.[15] Furthermore, their cell wall-based mode of action, and the countless AMP motifs they present make bacterial resistance less likely.[16] However, concern about toxicity in humans has hindered their development.[16] Human neutrophil defensin 1 active peptide (HNP-1) derived from the innate immune system resolves this issue.[17] It is effective against a broad range of microorganisms such as E. coli with a preference towards Gram-positive bacteria including S. aureus, while also having low toxicity against mammalian cells, demonstrated in in vivo studies against M. tuberculosis.[18–21]

Aside from attachments or coatings, the suture can be improved as well with novel polymers, notably spider silk, the strongest known biomaterial.[22, 23] The synthesis of multifunctional silk using specific silk glands to assist in reproduction and predation is a unique ability of spiders following more than 380 million years of evolution.[24, 25] Spiders produce a silk with high tensile strength, temperature resilience, and bio-compatibility that makes it a superior material for medical sutures.[26–29] Spider silk proteins (spidroins) (Fig. 1A) consist of nonrepetitive N and C terminals shown to aid silk formation, and extensive repetitive regions in between that contribute to the properties of the silk.[30–36] For example, major ampullate spidroin has poly-A motifs in the repetitive region (Rep) that forms into crosslinked beta-sheets, giving the silk incredible strength.[37, 38]

**Figure 1.**
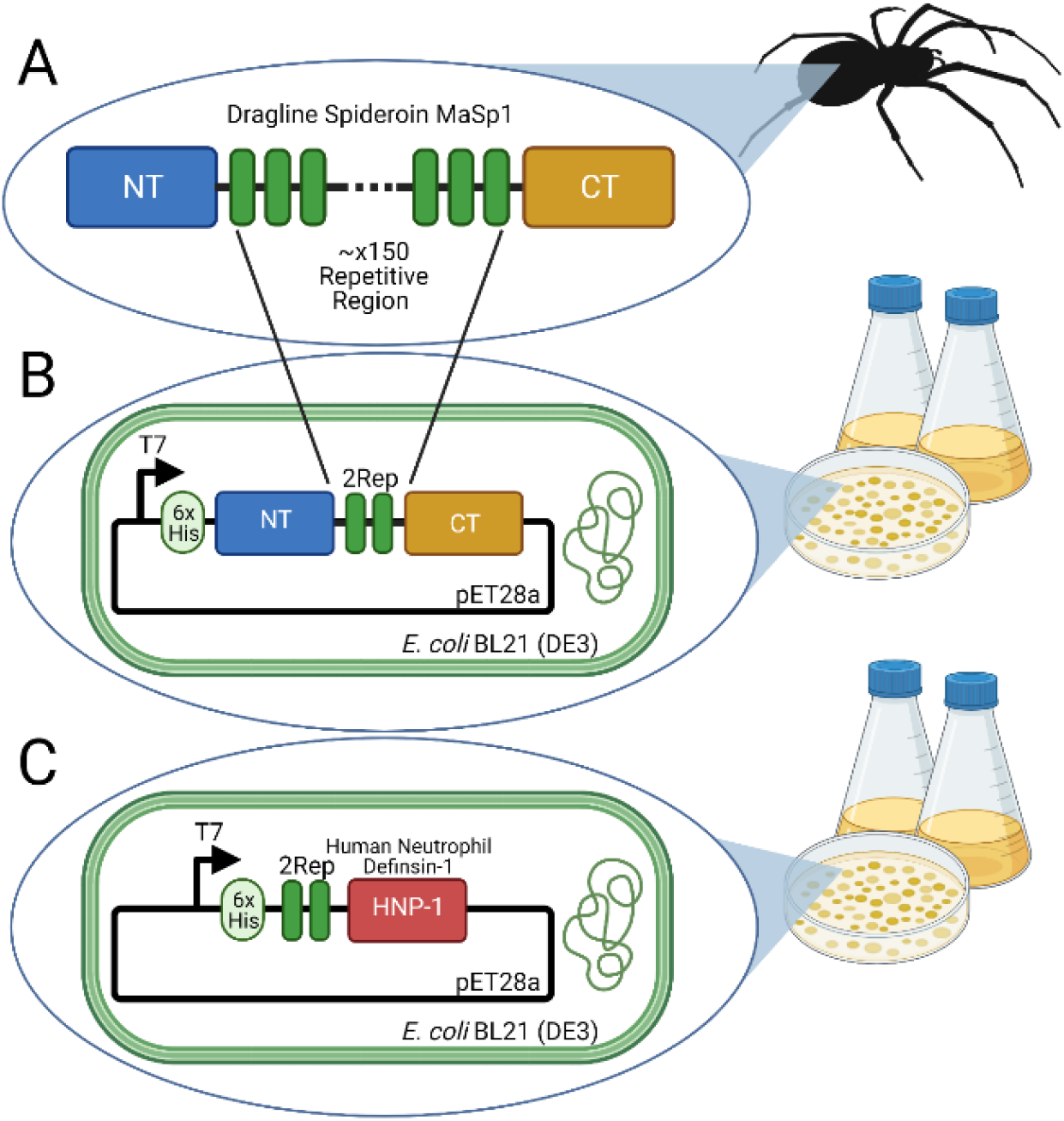
Recombinant spidroin design. **A)** Spidroins have a large repetitive region, most notably with the Poly-A motif. **B)** Mini-spidroin NT2RepCT reduces the repetitive region and adds a 6xHis tag for purification. The chimeric NT and CT increase solubility. **C)** 2Rep-HNP1 fusion peptide in same construct 6mer-HNP1 (mer is another repeat) maintains broad range antimicrobial activity after a coagulating bath. It was designed as a post-spinning coating for Perma-Hand sutures, and cannot take advantage of the unique strength and extensibility spider silk offers. Furthermore, the high molecular weight of this toxic protein decreases yield. [10, 11]

With E. coli protein expression systems growing in popularity, it is now cost-effective to produce recombinant spidroins. Smaller recombinant spidroins (minispidroins) with a reduced repetitive region can have much higher protein yields without compromising strength.[39] The chimeric minispidroin NT2RepCT (Fig. 1B) is a well-studied example that adds chimeric terminals with extreme solubility to increase yield further, up to 125mg/L.[35] To biomimetically spin NT2RepCT, a syringe pump pushes NT2RepCT into a coagulating bath (pH=5, isopropanol, or methanol) with sufficient shear force, which is then collected on a reel (Fig. 3A). NT dimerization stabilizes the fiber while CT amyloid-like fibril formation triggers solidification of the repetitive region (Fig. 2). [35, 36] This method of silk spinning has been shown to preserve beta-sheet formation and crosslinking tendencies of spidroins: 2Rep-sfGFP mixed NT2RepCT before spinning will attach to the main silk, tagging sfGFP while reinforcing the fiber.[36] This suggests that the nRep-xx fusion protein could potentially be used to tag peptides to recombinant spider silk.[36] HNP-1 is an impactful candidate.

**Figure 2.**
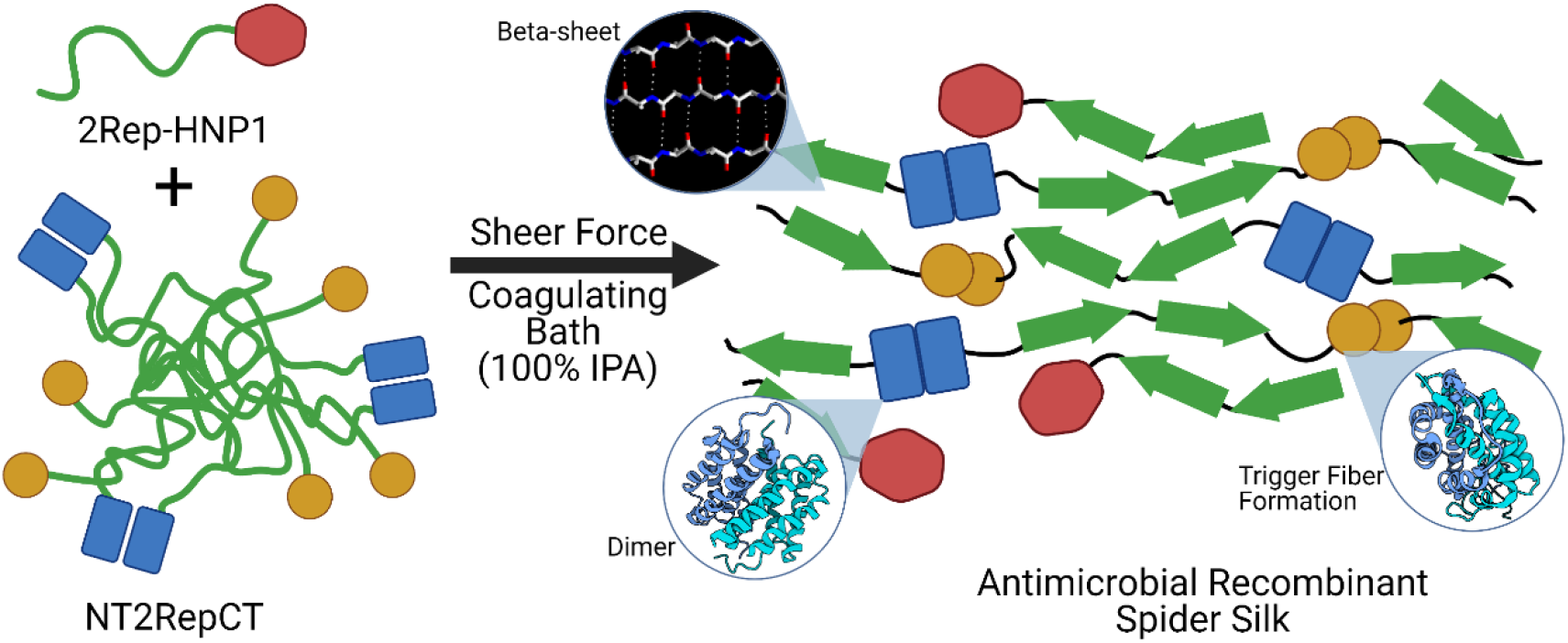
Mechanism of 2Rep-HNP1 attachment and silk formation. In native state, NT2RepCT micelles can maintain solubility at high concentrations because of the chimeric soluble NT and CT. After passing through a 100% IPA bath with sheer force, NT dimerization helps connect separate NT2RepCT together, and CT amyloid-like fibril formation triggers silk formation, turning the repetitive region into antiparallel bonding beta-sheets. In a mixture of containing 2Rep-HNP1, beta-sheet hydrogen bonding between the 2Rep domains of 2Rep-HNP1 and NT2RepCT can connect the two together and create antimicrobial recombinant spider silk to be used in sutures.

**Figure 3.**
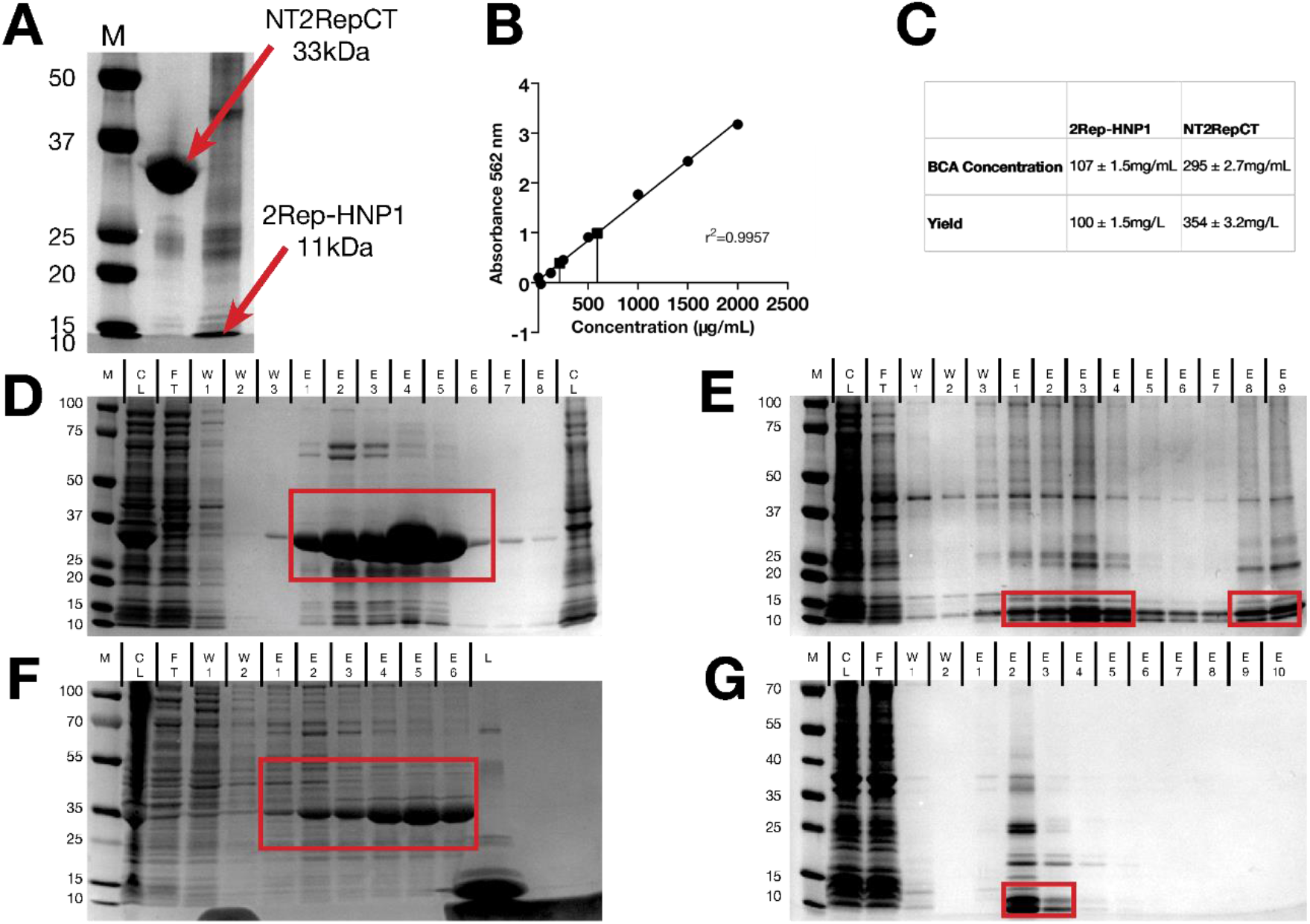
Ni-NTA protein purification results. **A)** Confirmation of 3k MWCO dialyzed protein. **B)** BCA curve with interpolated 1/500 dilution concentrations. **C)** Protein concentration and yield. **D, E)** Modified protocol purification results for NT2RepCT and 2Rep-HNP1. **F, G)** Original protocol results with low yield. Boxed fractions were used in dialysis. **A** is 12% Tris-Gly, **DEFG** are 8-16% Tris-Gly. M for **ADE** is Bio-Rad Dual Color, **F** is Thermo PageRuler, **G** is Thermo PageRuler Plus. **F** lane “L” is 10kDa lysozyme.

This study investigates whether the novel fusion protein 2Rep-HNP1 (Fig. 1C) can be produced and spun with the chimeric minispidroin NT2RepCT to create a silk product that paves the road for an antimicrobial spider silk suture that reduces surgical site infections. If 2Rep-HNP1 with a growth inhibiting effect on S. aureus and E. coli is mixed with NT2RepCT and biomimetically spun into silk, the 2Rep domains will form beta-sheet linkages and create a stable antimicrobial recombinant spider silk (Fig. 2). Furthermore, efficiency of spidroin production, adjustable silk diameter, and spidroin “sheet” formation were investigated during experimentation to create a cohesive demonstration of the practicality and far-reaching potential of antimicrobial recombinant spider silk.

## 3. Materials & Methods

### 3.1. Design and synthesis of chimeric spidroins

The chimeric minispidroin NT2RepCT (BBa_K3264000) is made of the N-terminal of the E. australis MaSp1 (AM259067), two repeats of the E. australis MaSp1 repetitive region (AJ973155), and the C-terminal of the A. ventricosus MiSp1 (JX513956).[35] The fusion protein 2Rep-HNP1 is made of two repeats of the E. australis MaSp1 repetitive region (AJ973155), and the Human Neutrophil Defensin-1 active peptide (P59665.1).[10] All segments are joined by GNS linkers, and a 6xHis (MGHHHHHH) tag was placed before the N terminal.[36] E. coli protein expression codon optimization was performed with GenSmart. Genes were synthesized and cloned into pET28a(+) vector by Genscript, and resuspended to 0.2μg/μL with ddH2O.

### 3.2. Bacterial transformation and expression

50μL aliquots of E. coli BL21(DE3) (Thermo Fisher) were mixed with 3μL of each plasmid with ddH2O as a control. Samples were incubated on ice for 30 min, 42C for 30 sec, and ice for 2 min. 250μL of S.O.C. media was added to each sample and shaker incubated (New Brunswick, Eppendorf) at 37C 225rpm 1 hr. 50μL and 200μL aliquots were spread on LB-kanamycin (50μL/mL) agar plates and incubated overnight at 37C.

Inoculated starter cultures of 6mL LB-kanamycin that were shaker incubated at 37C 250rpm overnight were used to make 25% glycerol stocks and inoculate in a 1:50 ratio TB-kanamycin in baffled 1L flasks (Thermo Fisher). Cultures were shaker incubated at 37C 250rpm until OD600 (Nanodrop, Thermo Fisher) was 0.6-0.8. NT2RepCT cultures were induced to 0.5mM IPTG, at 25C 250rpm overnight. 2Rep-HNP1 cultures were induced to 1mM IPTG, at 37C 250rpm 4hrs.

Bacteria pellets were harvested by ultracentrifugation (Sorvall, Thermo Fisher) at 4C 7000xG 15min, and twice washed with ice-cold PBS. NT2RepCT pellet was frozen −20C overnight, and 2Rep-HNP1 pellet was resuspended in 1:10 culture volume of denaturing lysis buffer (100mM NaH2PO4, 10mM Tris HCl pH8, 8M urea, 1mM PMSF, 10mM imidazole, adjusted pH8.0) 4C overnight.[11] NT2RepCT was resuspended in 1:10 culture volume of native lysis buffer (20mM Tris HCl pH8, 500mM NaCl, 1mM PMSF, 10mM imidazole, 100ug/mL lysozyme) and incubated 4C for 30min. Both were sonicated at 40% power 6 x 30sec and ultracentrifuged at 4C 13000xG 30min to recover lysate supernatant. Additional sonication and ultracentrifugation were performed if lysate was turbid.

### 3.3. Ni-NTA affinity chromatography purification

Native buffers in PBS with varying imidazole were prepared for NT2RepCT: equilibration (10mM), wash (20mM), elution 1 (100mM), elution 2 (250mM). Denaturing buffers in 100mM NaH2PO4, 10mM Tris HCl pH8, 8M urea with varying imidazole and adjusted pH were prepared for 2Rep-HNP1: equilibration (10mM, pH8), wash (20mM, pH6.3), elution 1 (100mM, pH5.8), elution 2 (250mM, pH4,5).

Ni-NTA column resin (HisPur, Thermo Fisher) equilibrated with two resin-beds of equilibration buffer was added 1:20 lysate and end-over-end shaker (Thermo Fisher) incubated at 4C overnight. The mixture was added to chromatography column (Pierce, Thermo Fisher) and passed flow-through twice. Columns were washed with 20 resin-beds of equilibration buffer and 10 resin-beds of wash buffer. Samples were eluted in 1 resin-bed fractions with 3 resin-beds of elution 1 buffer and 7 resin-beds of elution 2 buffer. SDS-PAGE of column fractions confirmed the protein of interest, using Tris-Gly 8-16% gels (Novex, Thermo Fisher) at 225V for 36 minutes stained with Blazin Blue (Goldbio) and imaged with FluorChem R (ProteinSimple).

The 6 most concentrated fractions were concentrated and desalted with 6mL 3MWCO protein concentrator columns (Pierce, Thermo Fisher) at 4C 4000xg 4 hrs and stored at 4C. BCA working reagent (Pierce, Thermo Fisher) with a 1:20 BSA 2000, 1000, 500, 250, 125, 25, 0 ug/mL standard curve and 1/500 and 1/1000 dilutions of concentrated protein in replicate was incubated at 37C for 30 minutes and measured at 562nm with Nanodrop One to find protein concentration and yield in mg/L culture.

### 3.4. Biomimetic silk spinning

NT2RepCT was diluted to 150mg/mL and 2Rep-HNP1 to 100mg/mL with 20mM Tris HCl pH8. Mixtures of 10%, 5%, 4% 2Rep-HNP1 with NT2RepCT and pure NT2RepCT were tested at 10-30μL/min with a 1mL (BD) syringe pump (Braintree Scientific) and 26G needle (BD)

The vertical pump ejected protein into a 100% isopropanol coagulating bath, and tweezers carefully collected continuous fibers and acted as a collection frame. Dried fibers were washed twice in ddH2O and stored at room temperature in petri dishes sealed with tape. Diameter measured after drying.

### 3.5. Silk diameter characterization

Prepared silk samples either in petri dish or on cardboard frame were viewed with the ZOE Fluorescent Cell Imager (Bio-Rad) in the brightfield and green channels. Random selections were used to calculate the mean diameters of the silk with ImageJ. Performed with 12 duplicates per silk.

### 3.6. Variable needle and rate spinning

30μL samples of NT2RepCT were spun at 15μL/min with 24G, 26G, and 28G needles. Diameter measured after drying. 30μL sample of NT2RepCT was spun with rate increasing to 150μL/min with 26G needle. After drying, silk sheet was carefully torn apart to reveal internal structure.

### 3.7. Silk degradation characterization

Silk samples are too light to quantify with a milligram balance. Quantitative criteria for degradation were established: Intactness, diameter, overall size, comparison to control once dry. Control was dry room-temperature silk. All samples were washed twice with ddH2O and dried before testing.

Physiological conditions were mimicked with PBS pH7.4. Sterilization was mimicked with 70% ethanol. Samples were photographed, and incubated in 1mL solution at 37C for 3 days monitored daily. Dry-cycle steam autoclave 121C 20min was also used to mimic sterilization.

### 3.8. Log reduction of S. aureus and E. coli CFU

Antimicrobial ability of 2Rep-HNP1 was assessed with E. coli BL21(DE3) and S. aureus Rosenbach 6538 ATCC. [Note: 6538 is NOT a MRSA strain.] 6mL LB cultures were shaker incubated 37C 150rpm overnight, pelleted at 4C 10000xg 4 min, and washed twice with ice-cold PBS. Bacteria was resuspended in ice-cold PBS to a final OD600=0.3 measured with Nanodrop One. The OD600=0.3 suspensions were diluted to OD600=0.1 with PBS and 1:1 added to 50μL samples of 2Rep-HNP1 (50, 25mg/mL), and shaker incubated at 37C 150rpm for 24 hrs. 50μL serial dilutions (1/10, 1/100, 1/10000, 1/100000) into PBS were plated on LB agar and incubated at 37C for 24 hours. ddH2O and NT2RepCT were controls. Performed in duplicate. Log reduction of CFU was calculated using OpenCFU to count colonies.[40]

### 3.9. Radial diffusion assay

50uL of E. coli and S. aureus OD600=0.3 suspension was plated in LB Agar with a spreader. 40uL of 2Rep-HNP1 (1mg/mL) was put on plates and incubated at 37C for24 hrs. NT2RepCT was used as control. Performed in duplicate.

### 3.10. Analysis and figures

All data are given as mean ± standard deviation unless noted otherwise. For analysis, one-way ANOVA test and Tukey’s multiple comparison test were performed, with p<0.05. Graphs were and plotted using GraphPad Prism 9.0. Schematics were made with BioRender.

## 4. Data & Results

### 4.1. Protein production with altered protocol

The final expression protocol used Terrific Broth expression media with 0.4% glycerol and 1:5 filled baffled 1L flasks. The final NT2RepCT native lysis buffer contained NaCl, a nonspecific Ni-NTA binding reducer, PMSF, a protease inhibitor, and lysozyme, a supplement to sonication. These additions were not harmful after desalting through 3k MWCO protein concentrator (Fig. 3A). BCA assay (Fig. 3B) calculated a concentration of 295±2.7mg/mL and final yield of 354±3.2 mg/L of TB culture (Fig. 3C). This value is statistically significantly higher than the original publication 125mg/L and the hitherto highest reported yield of 336nm/L.[35, 36]

The 2Rep-HNP1 protocol was modified to denaturing lysis buffer and a shorter induction time of 4hrs. BCA assay calculated a concentration of 107±1.5mg/mL and final yield of 100±1.5 mg/L of TB culture. After 7 runs final Ni-NTA purification with modified protocols (Fig. 3D, 3E) was much more efficient than original run (Fig. 3F, G).

After desalting, NT2RepCT and 2Rep-HNP1 both did not precipitate at high concentrations, and did not degrade for at least 1 week stored at 4C and with constant use at room temperature. Dilution to working concentrations of 150mg/mL and 100mg/mL for NT2RepCT and 2Rep-HNP1 with 20mM Tris did not lead to precipitation either.

### 4.2. Silk spinning with variable conditions

A simplified silk spinning device (Fig. 4A) was set up as described, with a bath of 100% isopropanol, syringe pump set to set to 15μg/min and with a 28G needle. Testing with only NT2RepCT revealed fiber formation was only possible when the needle was 5cm or from the container bottom (Fig. 4B), as solid silk must form during the short fall so it can be collected by tweezers (Fig. 5A).

**Figure 4.**
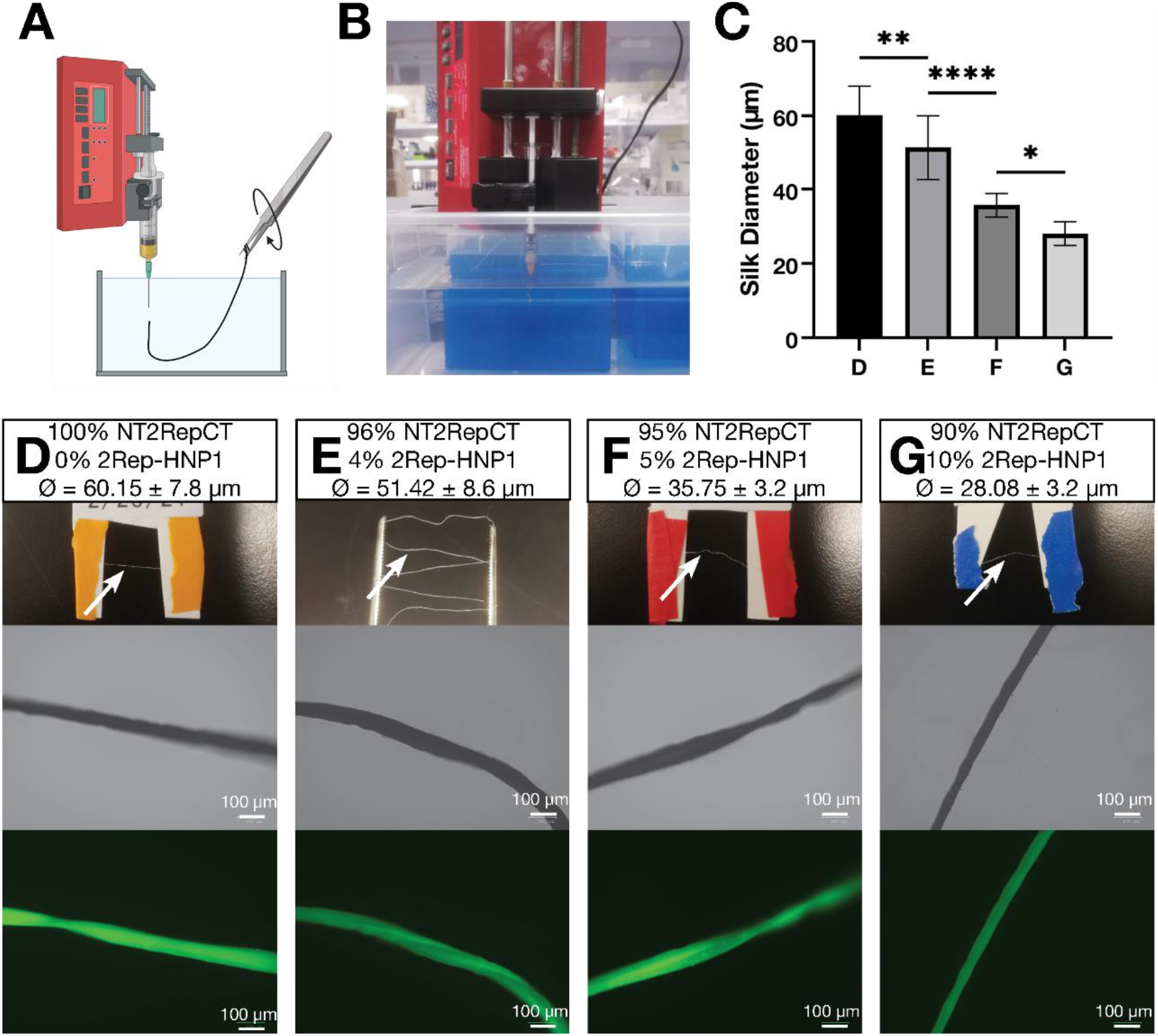
Biomimetic spinning of spider silk. **A)** Silk spinning components: a syringe pump, a coagulating bath, and a collection rack. **B)** Equipment setup and minimum container depth. **C)** Statistically significant diameter differences of mixtures demonstrate successful beta-sheet bonding. **D, E, F, G)** Normal and brightfield/green channel

**Figure 5.**
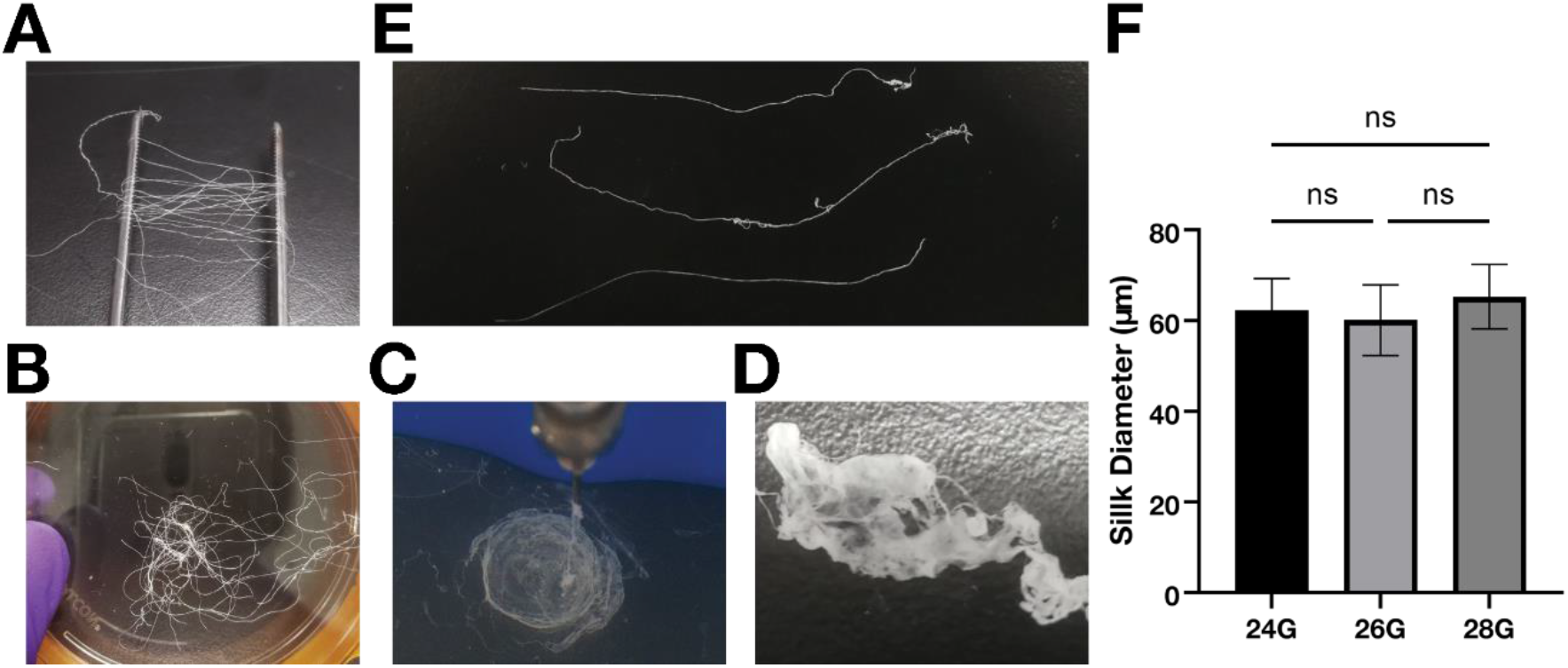
Supporting findings of silk spinning. **A)** Tweezer spinning frame. **B)** NT2RepCT silk in larger quantity. **C)** Circular sheet formed by rapid ejection. **D)** Dried sheet retains silk-like fibers. **E)** 4% silk degradation testing. Silks are control, second PBS 37C for 3 days, 70% EtOH 37C for 3 days. **F)** Needle diameter creates no significant difference in silk diameter.

Pure NT2RepCT (Fig. 4D, Fig.5B) had a mean diameter 60.15±7.8μm. Mixtures of 2Rep-HNP1 and NT2RepCT were tried. 10% 2Rep-HNP1 (Fig. 4G) partially precipitated when mixed with NT2RepCT and was difficult to draw into the syringe. Silk was discontinuous, breaking after contact with the container bottom, or upon attempting to gather a longer continuous segment. The longest collectable length was 3.81 cm. The mean diameter was 28.08±3.2 μm. Mixtures of 4% (Fig. 4E) and 5% (Fig. 4F) 2Rep-HNP1 had spinning properties similar to 100% NT2RepCT, with 4% less prone to breakage. The mean diameter of 4% silk was 51.42±8.6μm and the mean diameter of 5% silk was 35.75±3.2μm.

The statistically significant inverse relationship between 2Rep-HNP1 percentage and silk diameter (Fig. 4F) is accounted for by bonding of 2Rep-HNP1 to NT2RepCT denser crosslinking. [33, 36, 39] This demonstrates successful beta-sheet crosslinking. Explored more in discussion.

Pure NT2RepCT spun with a 26G (260μm) needle had a mean diameter of 60.15±7.8 μm when ejected at a rate of 15μL/min. Diameter did not significantly change with a 28G (184μm) or 24G (311μm) needle (Fig. 5C). This supports previous research findings between 32G and 34G needle use and demonstrates no correlation between silk diameter and needle diameter given a constant rate.[36]

Above 70uL/min, the syringe pump produced flat, circular sheets of solidified silk protein with a visibly uniform distribution (Fig. 5D). Pulling apart dried sheet revealed retention of fibrous properties (Fig. 5E), akin to tissues and other fiber-based materials.

### 4.3. Silk degradation conditions modeled in vitro

NT2RepCT and 4% 2Rep-HNP1 were incubated 37C for 3 days in PBS, 70% EtOH, or 20min in dry-cycle autoclave. After samples dried again, they were indistinguishable from the control according to criteria (Fig. 5E) and changes in silk diameter statistically insignificant.

### 4.4. S. aureus, E. coli log reduction by 2Rep-HNP1

S. aureus Rosenbach 6538 ATCC and E. coli BL21 (DE3) log reduction ability of 2Rep-HNP1 was tested at 50mg/mL and 25mg/mL, with no statistically significant difference in activity found. NT2RepCT had no effect on CFU log reduction, and has statistically significant difference in activity with 2Rep-HNP1 (Fig. 6A). Overall, for S. aureus 50mg/mL and 25mg/mL of 2Rep-HNP1 (Fig. 6B) had a 1.4-log and 1.2-log reduction, and 50mg/mL and 25mg/mL of NT2RepCT had a 0.05-log and 0.02-log reduction. For E. coli 50mg/mL and 25mg/mL of 2Rep-HNP1 (Fig. 6C) had a 2.1-log and 1.6-log reduction, and 50mg/mL and 25mg/mL of NT2RepCT had a −0.4-log and 0.07-log reduction. Control is 0 (Fig. 6 C, E). Log reduction demonstrates in liquid culture the broad-range antimicrobial activity of 2Rep-HNP1 against S. aureus and E. coli.

**Figure 6.**
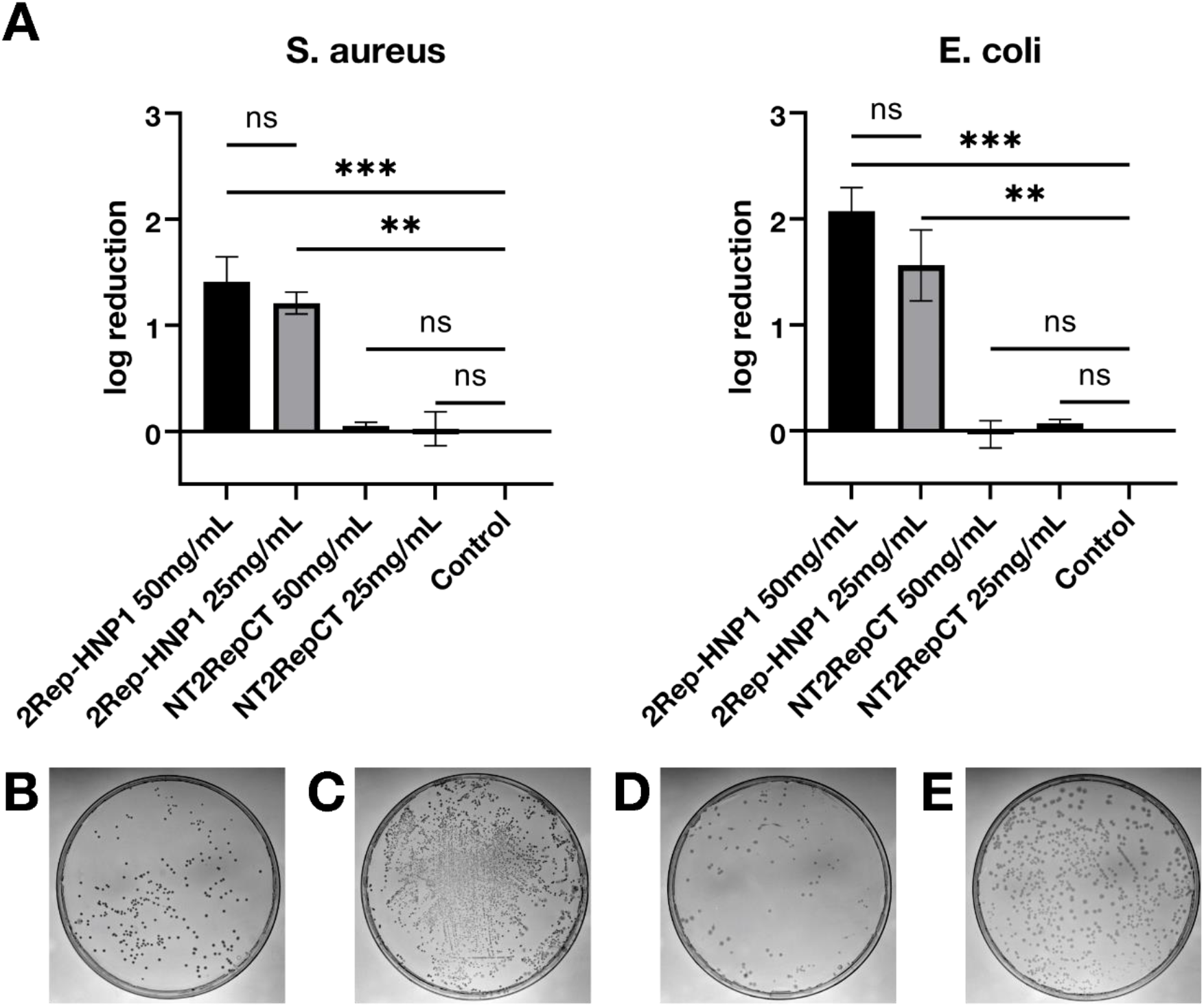
Antimicrobial testing of 2Rep-HNP1 with log reduction assay. **A)** Log reduction of CFU with 2Rep-HNP1. Controls represent inoculated media diluted with ddH2O instead of protein. NT2RepCT are a second control so peptide presence is not confounding. 50mg/mL and 25mg/mL did not produce a statistically significant in log reduction, but all samples reduced CFU at least tenfold. Log reductions were calculated with log (CFUControl/CFU). **B)** CFU count in S. aureus treated with 2Rep-HNP1. **C)** Control for S. aureus. **D)** CFU count in E. coli treated with 2Rep-HNP1. **E)** Control for E. coli.

30μL of 1mg/mL 2Rep-HNP1 was dropped on spread plates of S. aureus and E. coli formed semi-visible zones of inhibition, further demonstrating the antimicrobial activity of 2Rep-HNP1 in various conditions.

## 5. Discussion

Surgical site infections (SSI) represent a rising global health problem. Many infections come from surgical sutures, so antibiotic coated sutures have been developed in response. However, these hasten the antibiotic crisis and create harder to treat more lethal variants such as MRSA. Therefore, alternatives to antibiotic coated sutures must be researched.[5, 9–11] 2Rep-HNP1 beta-sheet crosslinked to NT2RepCT combines the antimicrobial peptide properties of Human Neutrophil Defensin1 with the strength and elasticity that spider silk offers.[10, 14, 33, 36]

To validate the main hypothesis and demonstrate the feasibility of this approach, three important criteria were met: antimicrobial activity, beta-sheet crosslinking, and stability.

Direct antimicrobial silk testing requires 3m of unwoven silk per replicate, which is only possible in industrial or highly specialized settings.[35] Log reduction of liquid culture is known to be a good approximate for antimicrobial activity after silk spinning, which makes it more suitable for small-scale preliminary investigation.[10, 11] Results demonstrate the antimicrobial activity of 2Rep-HNP1 against E. coli or S. aureus is not affected by the addition of 2Rep (Fig. 1C, Fig. 6), with reduction of at least 90% in both. This is consistent with antimicrobial 6mer-HNP1, and the fluorescent 2Rep-sfGFP, and supports that Rep does not interfere with protein functions in pre- and post-spinning.[11, 36] The broad-range action of 2Rep-HNP1 against both Gram-negative and Gram-positive bacteria further demonstrates viability in sutures.

The inverse relationship between 2Rep-HNP1 percentage and silk diameter (Fig. 4C) confirms 2Rep beta-sheet crosslinking behavior (Fig. 2). Adding 2Rep-HNP1 to NT2RepCT causes crosslinking and creates creates a denser silk that may also be stronger, but overly high concentrations such as 10%, leads to precipitation and a discontinuous silk.[39] 4% silk is a good balance, being the most akin to pure NT2RepCT in terms of spinning properties and continuity, but is diameter difference shows 2Rep-HNP1 crosslinking still occurs. This shows modified 2Rep proteins can bind to NT2RepCT and be spun into silk using a simple syringe pump.

Sutures experience a wide range of clinical conditions, and must not degrade at any point. NT2RepCT, 2Rep-HNP1, and all spidroin proteins depends on beta-sheet crosslinking and crystallinity, and a critical failure could endanger patients.[40] Both NT2RepCT and 4% silk was shown (Fig. 5E)to not degrade at physiological temperature 37C while suspended in physiological pH buffer, in 70% ethanol, or when autoclaved. These conditions model real-world use, and demonstrates spider silk has the stability required of suture products.

These criteria validate the hypothesis: The novel 2Rep-HNP1, with growth inhibiting effect on S. aureus and E. coli, can attach via 2Rep beta-sheet linkage to NT2RepCT and be spun into silk that is stable in surgery settings. This silk is a solution to SSI due to sutures. Other testing rounds out this model: efficient silk production for use at industrial scales, and alternate methods of silk formation and use.

Previous production methods for NT2RepCT have used LB and a simple lysis buffer, to a published yield of 125mg/L.[35] The well-designed chimeric terminals allow for extreme solubility, so higher yield methods are worth exploring. Production with TB and a more complex lysis buffer increases yield to 354±mg/L (Fig. 3). This means 2.8km of silk can be spun with 1L of culture, a very efficient yield.[35] Ability of NT2RepCT and 2Rep-HNP1 to be stored at high concentration without special conditions lowers production costs. The high tolerance of needle diameter (Fig. 5F) for production of silk further lowers production cost. More efficient methods increase the likelihood of industrial production and product accessible, therefore real-world impact of antimicrobial recombinant spider silk.

Protein ejection rate alters the shape of the product into a “sheet”. This sheet still retains the fibrous nature of slower rate silk, but has a more random arrangement than woven silk and may be useful in situations were this is preferred, similar to paper structure. Sheets demonstrate how this protein combination can be adapted to any application where antimicrobial properties and the strength and flexibility of spider silk is needed, such as flexor tendon repair or liquid stiches.[41, 42]

## 6. Conclusion

2Rep-HNP1 and NT2RepCT form a stable antimicrobial spider silk material with potential use as sutures to solve surgical site infections. MRSA and other Gram-positive bacterial infections can be targeted by 2Rep-HNP1; this process does not disrupt HNP-1 function. Further testing provides a new protocol to increase yield of component proteins and supports the viability of industrial-scale production.

This provides evidence for a customizable, modular, functionalized spidroin 2Rep-xx. Not just antimicrobial peptides, any protein could potentially be added to NT2RepCT to create a strong silk with the properties of the added protein, or even of multiple mixed together. Further confirmation of the reliability of this method with other impactful proteins has the potential to usher in a new age of strong, elastic, spider silk-based biomaterials.

## 7. Acknowledgements

LabShares Newton and the Newton South High School Science Department provided lab space and materials. Snapgene provided software, Medsix provided the syringe pump, and BOA Biomedical provided S. aureus. S.L., H.S., and D.B. gave advice for science research. B.W., Y.Y.C, J.L. provided purification and silk spinning advice. L.B. and M.L. assisted in literature review of AMP uses. S.L. reviewed manuscript.

## 8. Attributions

F.Y.C.L. conceived the original idea, performed all lab work, performed all analysis, and made manuscript and figures.

The author declares no competing interests.

## Notes

### Competing Interest Statement

The authors have declared no competing interest.

